# An “individualist” model of an active genome in a developing embryo

**DOI:** 10.1101/2021.01.08.425929

**Authors:** Shao-Kuei Huang, Sayantan Dutta, Peter H. Whitney, Stanislav Y. Shvartsman, Christine A. Rushlow

## Abstract

The early *Drosophila* embryo provides unique experimental advantages for addressing fundamental questions of gene regulation at multiple levels of organization, from individual gene loci to the whole genome. Using *Drosophila* embryos undergoing the first wave of genome activation, we detected discrete “speckles” of RNA Polymerase II (Pol II), and showed that they overlap with transcribing loci. We characterized the spatial distribution of Pol II speckles and quantified how this distribution changes in the absence of the primary driver of *Drosophila* genome activation, the pioneer factor Zelda. Although the number and size of Pol II speckles were reduced, indicating that Zelda promotes Pol II speckle formation, we observed a uniform distribution of distances between active genes in the nuclei of both wildtype and *zelda* mutant embryos. This suggests that the topologically associated domains identified by Hi-C studies do little to spatially constrain groups of transcribed genes at this time. We provide evidence that linear genomic distance between transcribed genes is the primary determinant of measured physical distance between the active loci. Furthermore, we show active genes can have distinct Pol II pools even if the active loci are in close proximity. In contrast to the emerging model whereby active genes are clustered to facilitate co-regulation and sharing of transcriptional resources, our data support an “individualist” model of gene control at early genome activation in *Drosophila*. This model is in contrast to a “collectivist” model where active genes are spatially clustered and share transcriptional resources, motivating rigorous tests of both models in other experimental systems.

## Results and Discussion

Genome activation in *Drosophila* begins one hour after fertilization with the transcription of about 120 genes [1–3]. This limited number of active genes, together with the simple organization of syncytial nuclei in the early *Drosophil*a embryo, provided a unique opportunity to globally visualize RNA Polymerase II (Pol II) at sites of nascent transcription at the single-nucleus level, thus lending a complementary view to Pol II activity seen in ChIP profiles [2,3]. High resolution microscopy of embryos incubated with antibodies that recognize the RPB1 subunit of Pol II (see Methods) revealed distinct Pol II foci (Figure 1A), reminiscent of the membraneless condensates of Pol II seen at active genes in mammalian cells, also called “speckles” [4–9]. That the Pol II speckles were transcribing genomic loci was suggested by the observations that they appeared only after genome activation, being diffuse earlier (Figure S1A), and that they colocalized with fluorescent RNA probes for early transcribed genes such as *Bsg25D* and *slam* (Figure 1B).

**Figure 1.**
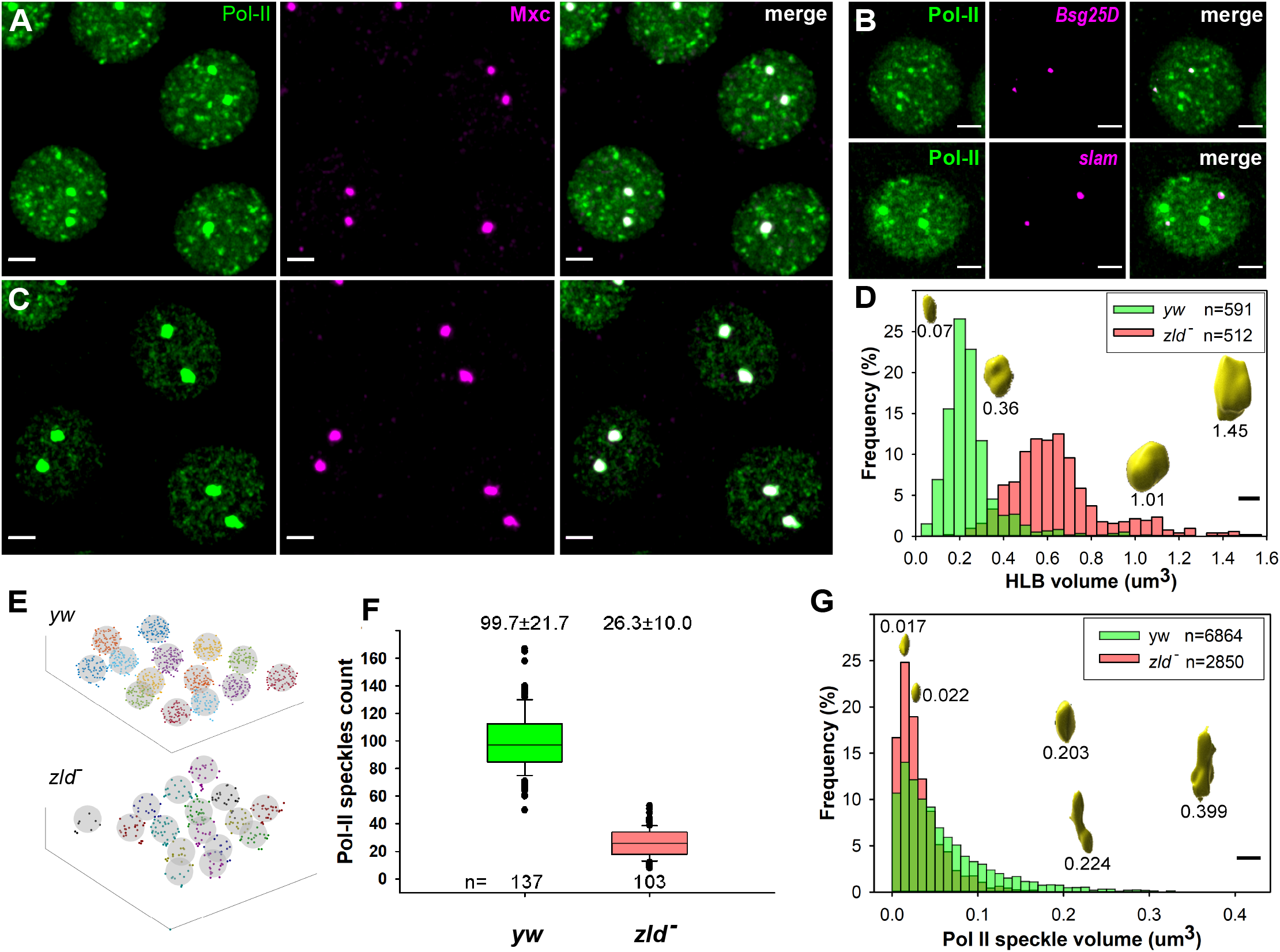
Pol II redistributes in the absence of Zelda. **(A and C)** immunofluorescence (IF) of wild type (*yw*, **A**) and *zelda* RNAi (*zld* ^*-*^, **C**) nc12 embryos with RNA polymerase II (Pol II) and Multisex Combs (Mxc) antibodies. Scale bar = 2 µm. **(B)** Pol II IF combined with RNA FISH of Zelda target genes, *Bsg25D* and *slam*. Scale bar = 2 µm. **(D)** HLB volume distribution in *yw* (green, 0.22±0.13 um^3^) and *zld* ^*-*^ (dark pink, 0.62±0.22 um^3^). HLB masks were generated by Imaris software; volumes are denoted next to masks. Scale bar = 1 µm. **(E)** Schematic diagrams of *yw* and *zld* ^*-*^ nuclei (gray circles) and the Pol II speckles allocated to them (dots). **(F)** Box plots of Pol II speckle counts in *yw* and *zld* ^*-*^ nuclei. **(G)** Volume distribution of Pol II speckles in *yw* (0.061±0.059 um^3^) and *zld* ^*-*^ (0.032±0.028 um^3^); speckle-mask volumes labeled. Scale bar = 1 µm.

Two of the Pol II speckles in each nucleus were always larger than others, and we speculated that they could be Cajal bodies or histone locus bodies (HLBs), multiprotein-RNA complexes involved in the transcription and processing of snRNPs and histone RNAs, respectively, that have been seen in *Drosophila, Xenopus*, and human cells [10–14]. We identified them to be HLBs, as they hybridized to H3 and H4 antisense probes (data not shown) and stained for Multi sex combs (Mxc), a protein component of the HLB [15–17] (Figure 1A and C, Mxc in magenta).

Since Zelda is a pioneer transcription factor involved in genome activation, and most early genes are highly downregulated in the absence of Zelda [1,18,19], we stained embryos devoid of maternal and zygotic *zelda* RNAs (Figure S1A), herein referred to as *zld*^*-*^ (see Methods [20]), for Pol II. As expected, there appeared to be fewer speckles in *zld*^*-*^ nuclei, but surprisingly, the HLBs appeared larger (Figure 1C; see volume sizes in Figure 1D), suggesting that in the absence of full genome activation, Pol II redistributes to the HLBs. In cases where only one HLB was observed, likely due to chromosome alignment [16], the one HLB had a volume larger than either HLB when two are present in a nucleus (Figure S1B).

To quantify the number and size of Pol II speckles per nucleus in our wild-type and *zld*^*-*^ nc12 embryos, each image with at least 14 nuclei, we used the Imaris “spot” finding function to call Pol II spots, and the Imaris ”surface” function to determine the center of each nucleus (see Methods). Figure 1E shows schematic 3D representations of wild-type and *zld*^*-*^ confocal images, with each spot assigned to the nearest nucleus (see Methods). Wild-type nuclei contained ~100 Pol II spots (99.7±21.7; n=37) versus ~25 in *zld*^*-*^ (26.3±10.0; n=103) (Figure 1F), and likewise, the global density of Pol II spots in *zld*^*-*^ embryos was about a quarter of that in *yw* (Figure S1D; see Methods). Importantly, the *zld*^*-*^ spots appeared smaller than those in *yw* (Figure 1G), consistent with the decreased transcriptional output observed for many patterning genes in *zld*^*-*^ [19,21–23]. To test this idea, we measured Pol II spot size at the *hunchback* (*hb*) locus, and observed smaller spot volumes in *zld*^*-*^ compared to *yw* (Figure S1E), in line with the reduced expression levels of *hb* seen in *zld*^*-*^ [19].

The observation that in the absence of Zelda, the HLBs become larger while Pol II speckles are smaller or disappear is reminiscent of a phenomenon that occurs with solid solutions in vitro known as Ostwald ripening [24], where smaller condensates shrink/dissolve and redeposit on to larger ones.

The ability to discern discrete Pol II speckles enabled us to examine the spatial organization of the early active genome. We speculated that the Pol II spots could be either uniformly distributed in the nucleus or somehow clustered. For example, there could be non-uniformity along the radius, i.e., a radial gradient, in one or both genotypes, or there could be local clustering (Figure 2A). First, we asked if there is a systematic inhomogeneity of density of spots as we scan radially outwards from the center of the nucleus. To address this question, we divided the nuclear regions into concentric 3D shells from the center of mass along the radius (like onion layers), with shell volumes getting smaller and smaller as they approach the center of the nucleus. Then we calculated the local density of spots for each shell, *ρ*(shell), i.e., the number of spots in each shell divided by the volume of the shell, and then divided by the average total nuclear density (*ρ*). Figure 2B shows that the *ρ*(shell)/*ρ* ratio for each shell along the radius was ~1, which indicates an even distribution of spots along the nuclear radius.

**Figure 2.**
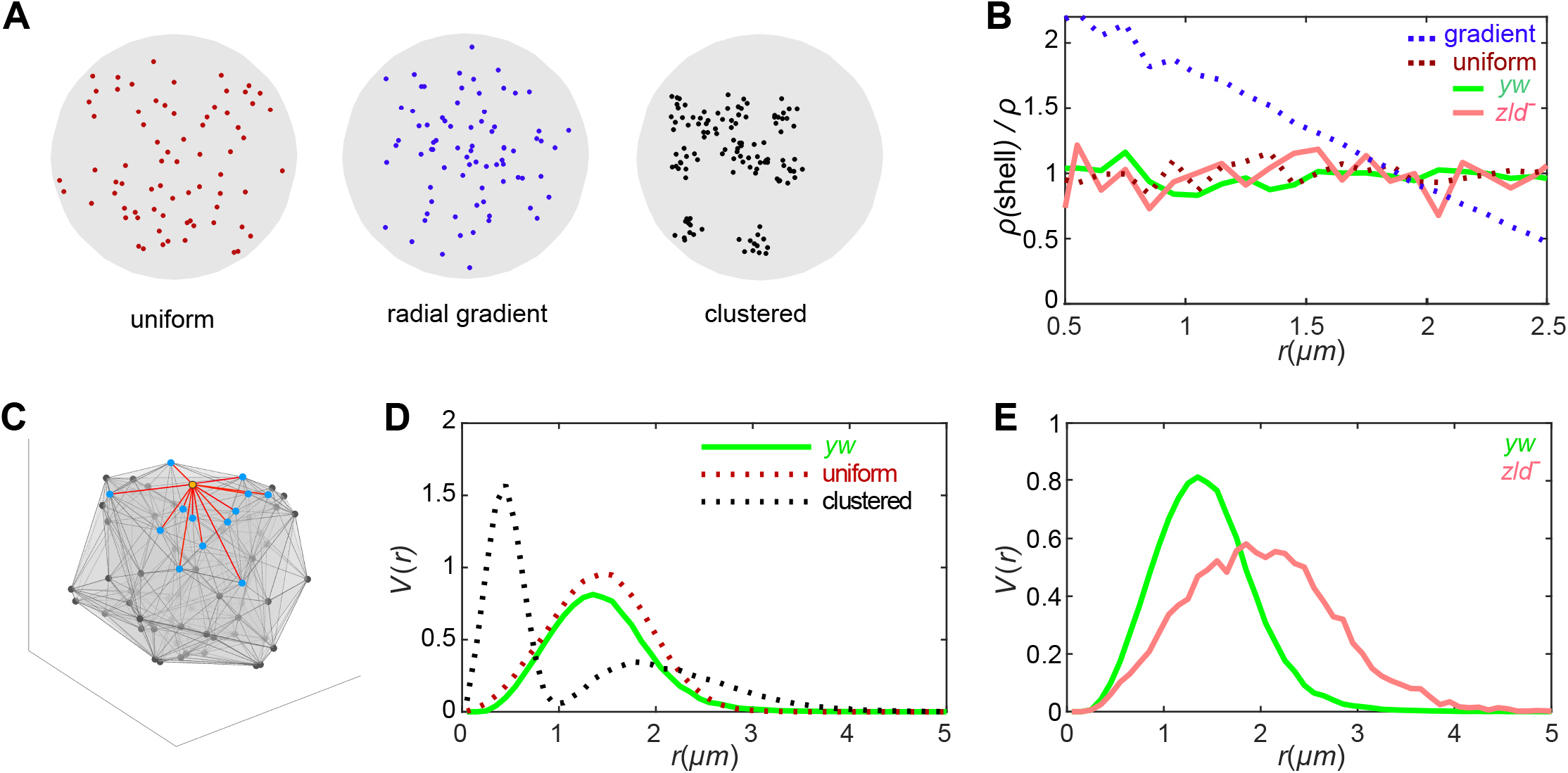
Pol II speckles are distributed evenly and randomly throughout the nucleus. **(A)** Synthetic point patterns distributed uniformly (left), in a gradient along the radius (center), or clustered within a nuclear sphere (right). **(B)** The ratio of shell density, *ρ*(shell), of Pol II spots to global density of spots, *ρ*, as a function of distance from the center of mass of each nucleus for *yw* (green) and *zld* ^*–*^ (dark pink) embryos. Hypothetical ratios are shown as dashed lines, uniform (red) and gradient (blue). **(C)** Schematic example of Pol II speckles in a nucleus. One speckle (orange), its neighbors (blue), and lines between them (red) are highlighted. **(D)** Probability distribution functions of the distance between nearest neighbors (*V*(*r*)) in *yw* (solid green line) compared to hypothetical uniform (red dashed line) and clustered (black dashed line) distributions in the same density shown in A. **(E)** Probability distribution function of distances between nearest neighbors (*V*(*r*)) in *yw* (green) and *zld* ^*–*^ (dark pink) nuclei. The shift of the dark pink curve to the right reflecting greater distances between spots is due to the reduced total number of spots in *zld* ^*–*^ embryos, while the shift down occurs since Y-axis measures frequency.

Next, we asked whether the Pol II speckles were locally clustered (Figure 2A) by assessing the distribution of the distances between spots and their neighbors. First, we took the 3D coordinates of the spots and constructed the Delaunay triangulation of the set of points using MATLAB (see Methods). Next, we identified all pairs of points in each tetrahedron to be each other’s nearest (Voronoi) neighbors [25] (Figure 2C shows a Delaunay tetrahedron with a spot and its neighbors highlighted in magenta), then calculated the probability distribution of distance between each pair of nearest neighbors *V*(*r*) (see Methods) [26]. Figure 2D shows these functions calculated for a uniformly dispersed (Poisson distribution of points) versus a clustered distribution of points generated in-silico as schematized in Figure 2A. If distributed uniformly, spot distances will fit a Gaussian distribution with a single peak (Figure 2D, red dashed line). If however spots are clustered, we would expect neighbor distances for spots within the same cluster to be small, while the distances between spots in different clusters to be large, therefore the distribution would have two peaks (Figure 2D, black dashed line). Wild-type spots fit a Gaussian distribution with one peak (Figure 2D, solid green line), as expected for a uniform distribution. Moreover, removing Zelda does not affect the distribution, but instead, the average distance between the neighbors is increased (Figure 2E, red curve shifts right), which is likely a consequence of the reduced number of Pol II speckles observed in *zld*^*-*^ nuclei (Figure 1F). These results show that Pol II speckles are uniformly distributed throughout the nucleus in both wild-type and *zld*^*-*^ nuclei, suggesting that although Zelda affects the localization of Pol II molecules in the nucleus, it does not change the 3D organization of the active genome.

Having quantified Pol II speckles in wild-type nc12 nuclei to be ~100, we wondered whether this number was consistent with the number of transcriptionally active genes estimated from genome-wide expression profiling and Pol II ChIP experiments. Liang et al. [1] reported that about 120 genes were down-regulated in *zld*^*-*^, and we obtained a similar number of genes highly bound by Pol II in nc12 embryos (see Methods) [3]. Could the lower number of Pol II speckles compared to active genes be due to multiple genes transcribing together in so-called Pol II factories, or hubs, where Pol II and associated co-factors concentrate together to increase transcriptional efficiency, firing, and output [9,27–34]? Whether transcriptional hubs are a general feature of co-regulated genes remains to be elucidated [35,36] and the nc12 embryo, in which most of the active genes are Zelda target genes (see Pol II and Zelda ChIP tracks in Figure S3A) presents a good system to address this question. Support for this idea comes from HiC studies of nc12 embryos that revealed the emergence of TADs during genome activation, as well as interactions between TAD boundaries [37]. Curiously, many of the nc12 TAD boundaries were positioned at active Zelda target loci, and several boundaries showed interactions (Figure S3C), which could occur in Pol II hubs.

Since it is not possible to know which gene or genes are being transcribed at a Pol II speckle, we instead looked for instances where transcribing genes come into close proximity in 3D, and then asked if they shared a Pol II speckle, i.e., Pol II hub. Focusing on a 5 megabase (Mb) region of chromosome 2L with several highly expressed genes (Figure 3A), we performed dual color RNA FISH of genes in pairwise combinations to first assess if their nascent transcription foci ever appeared close together despite lying far apart on the chromosome (see Methods) (Figure 3B-E). In each case, we observed the two FISH signal centers come within less than 400 nm of each other at least 10% of the time; 400 nm was the cutoff we used to define “close proximity”, as promoter-enhancer interactions have been observed at approximately this distance [38,39] (Figure 3H, dashed line demarcates 400 nm). For example, *CG15382* and *CG14014* are separated by 3.4 Mb, but their transcriptional foci were seen as close as 230 nm (Figure 3B).

**Figure 3.**
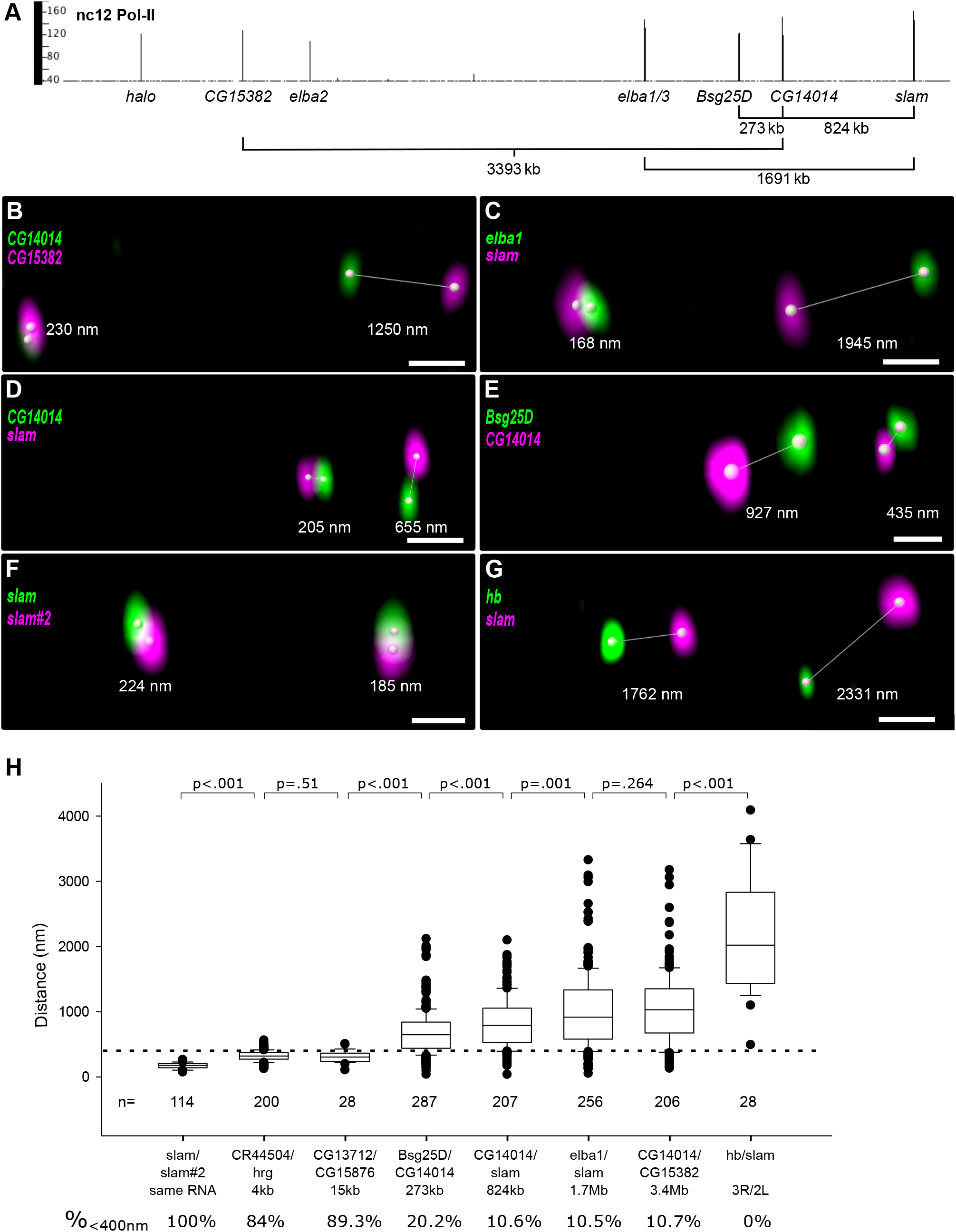
Distant loci occasionally transcribe in close proximity. **(A)** Pol II ChIP data [3] with one dimension (1D) distances labeled for several pairs of active genes in a 5 Mb region of chromosome 2L. **(B-G)** 3D views of single nuclei with dual color FISH of *CG15382* and *CG14014* (B), *elba1* and *slam* (C), *CG14014* and *slam* (D), *Bsg25D* and *CG14014* (E), *slam* and *slam#2* (F), and *hb* and *slam* (G). Distances are measured by Imaris software. Scale bar = 1 µm. *slam* and *slam* #2 are two probes against *slam* mRNA 78bp apart. **(H)** Box plots of 3D distances for each pair of genes. The pairs are ranked, left to right, by increasing 1D distance. 3D measurements (mean±SD) for each pair are: *slam/slam#2* (172±46 nm), *CG13712/CG15876* (309±87 nm), *Bsg25D/CG14014* (666±266 nm), *CG14014/slam* (825±370 nm), *CG14014/CG15382* (1327±791 nm), *hb/slam* (2186±909 nm). Note that the further away two loci are located on the chromosome in 1D, the greater the average 3D distance apart, up to a certain point, beyond which there was no longer a correlation, as no significant difference in the 3D distance distributions was observed between genes lying 1.7 Mb or 3.4 Mb apart (Mann-Whitney Rank Sum Test). This indicates that beyond a certain distance, chromosome movements in 3D are not predictable.

In a control experiment, two probes from the *slam* gene, 87 bp apart, showed distances in the range of 76 nm to 265 nm apart, with a mean of 172±46 (Figure 3F), thus we considered 218 nm the operational resolution limit in our imaging experiments. In contrast, two probes for genes on different chromosomes (*slam* on 2 and *hb* on 3) never came into close proximity (Figure 3G). Note that the closer two loci are in 1D, the more often their FISH foci centers came within 400 nm of each other in 3D (Figure 3H, bottom), however, beyond ~1 Mb, that number remained about the same, ~10%. Combining FISH with Pol II antibody staining, we asked whether two genes transcribing in close proximity could occupy a single Pol II speckle. Figure 4A shows images of *Bsg25D* and *CG14014* foci with their associated Pol II speckles, and in each case, when the two foci were between the resolution limit (~218 nm) but below ~400 nm, the Pol II speckles could be resolved as discrete spots (Figure 1G). Similar results were seen for FISH pairs, *slam* and *elba1* (Figure 4B), *slam* and *elba2* (Figure 4C), and *CG14014 and CG15382* (Figure 4D). A striking example of genes in close proximity was observed at the *elba1* locus where two sister chromatids were actively transcribing, each in its own discrete Pol II spot, which measured 223 nm apart (Figure 4E). These results indicate that genes in close proximity do not share Pol II hubs.

**Figure 4.**
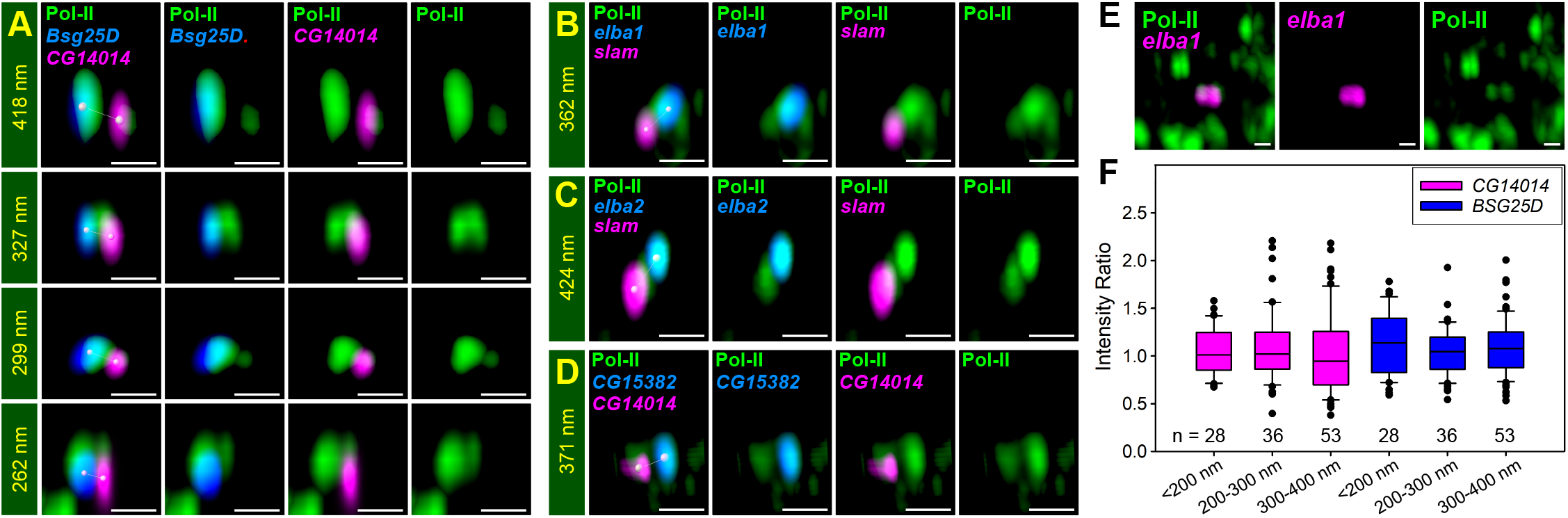
Zelda co-regulated genes do not share Pol II hubs or increased transcriptional output when in close proximity. **(A-D)** Zoom-in 3D images of Pol II IF (green) and dual color RNA FISH of *Bsg25D* (blue) and *CG14014* (magenta) **(A)**, *elba1* (blue) and *slam* (magenta) **(B)**, *elba2* (blue) and *slam* (magenta) (C), and *CG15382* (blue) and *CG14014* (magenta) **(D)** with 3D distances shown. Note that Pol II speckles overlap with each RNA FISH signal and are separable. **(E)** 3D images of Pol II IF (green) and *elba1* RNA FISH (magenta) on sister chromatids showing. Note the *elba1* signals have not separated completely, though the associated Pol II speckles are separable (233nm apart). **(F)** Distances of the pairs have no correlation with their expression level. The pairs are sorted into <200 nm, 200-300 nm, 300-400 nm, and >500 nm groups based on pair distance. Intensity ratio (Y-axis) is the smFISH signal of the indicated gene in the same nucleus, but sorted into different groups (X-axis). There is no significant difference between groups for both genes.

One possible explanation for the fewer number of Pol II speckles than transcribed genes is when genes lie within 1-2 kb of each other on the chromosome, for example, *elba 1* and *elba 3* (Figure 3A), their FISH and Pol II signals are too close to be distinguished (see Methods). This situation could give rise to a large or odd shaped Pol II speckle, which we observed at a low but consistent frequency in nc12 wild-type nuclei (see examples in Figure 1G). On the other hand the large speckles could comprise several distant genes transcribing together, as the evidence for TAD boundary interactions suggests [37].

Lastly, we asked whether the transcriptional output of genes transcribing in close proximity was increased compared to when they were not close, as this could be a consequence of sharing transcriptional machinery [34,40]. We used single molecule (sm) FISH (see Methods) to compare transcript levels of *Bsg25D* and *CG14014* when in close proximity (less than 400 nm) on one homolog versus not in close proximity (more than 500 nm apart) on the other homolog (Figure 4F). Comparing expression of the two gene pairs only when in the same nucleus reduces inter-nuclear differences in numbers of nascent transcripts. For each nucleus a ratio was generated by dividing the smFISH signals of the pair in close proximity (binned into three groups - less than 200 nm, 200-300 nm, and 300-400 nm) by the signal of the distant pair. If there was a higher transcriptional output associated with the pair in close proximity, the ratio would be higher than 1, however, in each bin, the ratio was ~1 (Figure 4F), indicating that there was no benefit in being transcribed in close proximity.

Taken together, our results demonstrate that the early Zelda co-regulated genes do not transcribe in shared Pol II factories/hubs even when within 400 nm of each other. Although we cannot rule out the possibility that on occasion two or more loci might share a Pol II hub, either at nc12 or later when many more genes become active, our results support an “individualist” model, whereby each active gene transcribes in its own discrete Pol II hub. The individualist model is in contrast to the ‘collectivist’ model where active genes are spatially clustered and share transcriptional resources, motivating rigorous tests of both models in other experimental systems.

## Acknowledgements

The authors would like to thank Robert Duronio for the Mxc antibody, Stephen Small for the *hb* RNA probe, and Sevinc Ercan for her always insightful ideas over the course of this study. The research was supported by National Institute of Health (NIH) research grants: RO1GM63024 to CAR, T32HD7520 Training Program in Developmental Genetics to PHW, and the following grants to SYS: DP2EB024247, F32GM119297, RO1HD0858870.

## Author Contributions

All authors contributed to the design of the study. SKH carried out the imaging experiments and image analysis, SD carried out the Pol II distribution analysis and modeling, and both prepared the figures. All authors contributed to writing the manuscript.

## Methods

### Contact for Reagent and Resource Sharing

Requests for further information, resources, and reagents should be directed to and will be fulfilled by the Lead Contact, Christine A. Rushlow.

## Experimental Model and Subject Details

All flies were grown on standard fly (*Drosophila melanogaster*) cornmeal-molasses-yeast media. *y*^*1*^*w*^*1118*^ flies were used as wild type. Embryos were depleted of maternal and early zygotic Zelda using the Maternal-Gal4-shRNA system[41], where *MTD-Gal4/UAS-shRNA-zld* females (Sun et al., 2015) were crossed to *y*^*1*^*w*^*1118*^ males; the resulting embryos were devoid of *zld* RNAs and referred to as *zld*^*-*^.

## Methods details

### immunofluorescence (IF)

Embryos were collected 1 to 2.5 hours after egg laying at room temperature, dechorionated with Clorox, fixed in 4% formaldehyde/heptane, and devitellinized with methanol. Fixed embryos were rehydrated and stained with primary antibodies overnight, washed, and then stained with secondary antibodies conjugated with Alexa Flour 488, 555, or 647 (Invitrogen and ThermoFisher Scientific) for 2 hours at room temperature. After Hoechst 33342 (Tocris Bioscience) staining for 15 minutes, embryos were mounted on microscope slides with Aqua-Poly/Mount (Polysciences) and Number 1.5 glass coverslips (Fisher Scientific). Primary antibodies included: anti-RBP1 conjugated with alexa488 (CTD4H8, Sigma-Aldrich, 05-623-AF488), which detects all forms of RNA polymerase II (Pol II; 1:100 dilution); guinea pig anti-Multi sex combs (Mxc) to detect the histone locus body, a kind gift from Dr. Robert J. Duronio (1:10000 dilution); rabbit anti-Zelda antibodies raised against the N-terminal domain of Zelda, amino acids 1-517 (Pocono Rabbit Farms).

### Pol II speckle distribution analysis

For the Pol II foci distribution analysis, embryos were stained with the IF protocol described above except the mounting media Aqua-Poly/Mount (Polysciences) was diluted to 70% with PBS/Tween to prevent embryo drifting. Embryos were re-suspended in mounting media and placed on a 35mm glass-bottom dish (Bioptechs) without a coverslip to minimize deformation of nuclei. Visually sorted nc12 embryos (six *yw* and five *zld*^*-*^) were imaged. Pol II speckles were called using the Imaris “spot” function with the method of standard deviation between center-to-surround signal, and the center radius was set at 330nm. The threshold was determined empirically to include focused spots of signal with intensity greater than diffuse-looking nuclear signal. Diffuse Pol II spots lying outside of the nucleus and large spots comprising the HLBs as well as the centrosomes were removed. The centers of nuclei were determined by the preconverted signal of Pol II flooded inside the nuclei (Figure S2E) via the Imaris “surface” feature to mark surfaces of nuclei (Figure S2F) and calculated the center of the homogeneous mass. The average diameter of a nucleus (4.76±0.25 m) was determined by measuring the surface of the nuclear envelope with Lamin antibody staining (Hybridoma Bank, ADL84.12; 1:300 dilution) with Imaris ”surface” function. The mean plus three standard deviation 5.51 m) was used in Figure 1E. To distribute the dots to individual nuclei, we found the nucleus whose center of mass was nearest to the dot.

Number density of Pol II spots: To calculate number density of Pol II spots (*ρ*), we drew an imaginary sphere of radius *D*_0_, the average diameter of nucleus (4.76 m) centered at center of mass of each nucleus, and counted the number of spots inside it and divided it by the volume of the sphere to obtain the number density of Pol II spots in each nucleus. The mean and standard deviation is calculated for all nuclei in wild type and the *zld*^*-*^ embryos. We averaged over 300 spheres for both cases and densities for wild type and *zld*^*-*^.

Radial distribution of density: To quantify how the density of spots varies with the distance from centers of mass of the nuclei, we drew spherical shells of thickness 0.1 m centered at the center of mass of each nucleus and calculated the number of Pol II spots in each shell. We defined *ρ*(*r*), local density at distance *r* from the center of mass as the ratio of the mean number of particles in the spherical shell at distance *r* (averaged over all wild type and *zld*^*-*^ embryos separately) and the volume of the spherical shell at distance *r*. Figure 2C suggests that local density at all distances from the center is close to the global density of dots in both wild type and *zld*^*-*^ nuclei.

Distribution of distances between neighbors: To check if there is any correlation between the position of Pol II spots within nuclei, we calculated *V*(*r*), the probability distribution function of distance between nearest neighbor spots. To find the nearest neighbor of each spot, we found the Delaunay triangulation of the point pattern (where each point represents a Pol II spot) using the *delaunayTriangulation* function in MATLAB. Next, we allocated all pairs of points in each Delaunay tetrahedron to be each other’s nearest neighbors. Then, we calculated the probability distribution of distance between each pair of nearest neighbors.

Generation of synthetic point patterns: To check whether the Pol II spots within the nucleus are uniformly distributed or clustered, we generated synthetic point patterns, uniformly distributed within a sphere and distributed in a few clusters, with number densities similar to Pol II spots in the wild-type and *zld*^*-*^ nuclei. To generate a uniform (Poisson) distribution of particles, we took a sphere of diameter *D* and generated *ρ D*^*3*^*/6* particles within the sphere using the random number generator in MATLAB. To generate clustered distribution of particles, we took a sphere of diameter *D* and divided *ρ D*^*3*^*/6* particles into *m* clusters. First, we placed *m* points within the sphere randomly as the center of each cluster. Next, we drew a sphere of diameter *f*(*D*/*m*)^⅓^, where f<1, around each center. Finally, we distributed *ρ D*^*3*^*/6m* particles randomly into each of the *m* clusters. For the illustration in Figure 2, we used f=0.25 and m=10.

### Volume quantification

The volumes of Pol II speckles and HLBs were calculated using the “surface” function in Imaris software. First, the background subtraction level was established to generate the spots mask to define the edges of all possible spots. Second, the thresholds for calling spots of Pol II and Mxc, an HLB marker, were adjusted to label strong signals using the “spot” function as described above. The volume was calculated in 3D from each spot-center to the edge of the mask. When two or more spots were in the same region of interest, the volume was split. Two or more Pol II spots may be recognized as one when Imaris did not call them as adjacent spots due to the spot intensity level or the resolution limit. There are ~10 Pol II spots per nucleus with a volume size greater than 0.264 m^3^ and could be this type of fusion spots.

### Fluorescent *in situ* hybridization (FISH)

Hybridization of fixed embryos was done following a standard RNA FISH protocol [42]. The template DNA fragments were generated by PCR with anti-sense cDNA primers also containing the T7 promoter sequence. DIG or Fluorescein-labeled (Roche) anti-sense probes of *CG15382, elba1, Bsg25D, CG14014, hunchback* (*hb*), *CG15876, CG13712, slam* and *slam*#2 (two different regions of the *slam* gene) were generated by *in vitro* transcription with RNA labeling kits (Roche). Alexa Flour 488, 555, or 647 conjugated secondary antibodies were used to detect DIG or Fluorescein (FL) antibodies. When two genes were compared in experiments combining IF and FISH, IF was done prior to procedure FISH.

### High resolution imaging acquisition

Images of IF, FISH, and smFISH were acquired with the Zeiss confocal microscope, LSM 880, using the 100x objective lens. For the distance-measuring experiments, fluorescent beads (FocalCheck thin-ring fluorescent microspheres kit; Molecular Probes) were used to ensure that lasers were aligned within 50nm tolerance. Confocal settings were: 1156×1156 pixel, 25-30 z-stacks (except for the Pol II distribution experiment, which was scanned in 70 z-stacks) 0.15µm apart, 8 bit. Airyscan captured images were processed with the Airyscan Processing feature in the Zeiss Zen black software. 3D images of Pol II speckle masks, HLB masks, nuclear masks, dual-FISH pairs, and Pol II with dual-FISH pairs were snapshots from Imaris 3D view.

### ChIP-seq tracks of Pol II and Zld and Hi-C contact map

The ChIP-seq tracks of Pol II [3] and Zelda [18] of a 2 MB region of chromosome 2L (shown in Figs. 2 and S2) were generated using the Integrated Genome Browser (version 9.1.4, https://www.bioviz.org/); Y-axis, normalized sequencing reads [3]. Genome-wide, we estimated the total number of highly bound genes to be 143 (using a cutoff of 30), 111 (using a cutoff of 40), or 83 (using a cutoff of 50). Figure 3A shows a Hi-C contact heatmap of the corresponding region at nc 12 was generated from Hi-C data [37] using Hi-glass software (https://higlass.io/).

### Dual-color FISH 3D distance analysis

After stacks of images were loaded into Imaris, the position of foci were determined with the Imaris “spot” function to detect up to 8 foci per nucleus. Duplicated sister chromatids were excluded from further analysis. Samples were mounted with coverslips to flatten embryos in a traditional manner so the measured distance may be larger than actual length. Distances between foci of different genes were measured with the Imaris “measure” function by assigning start and end points from the center of defined foci. Distances between FISH foci for two different genes were measured for the following pairs of genes (1D distances listed, starting with closest; also see Figure 3H): *slam* and *slam*#2 (*slam*-RC: 1500 to 3427 and 425 to 1412, respectively, 87 bp apart; *CR44504-RA* (+144 to +900) and *hiiragi*-RD (+1273 to +3038), 4 kb apart; *CG13712*-RA (−111 to +660) and *CG15876*-RA (+29 to +478), 14.4 kb apart; *BSG25D*-RD (+10 to +2380) and *CG14014*-RB (+58 to +1034), 272.5 kb apart; *CG14014*-RB (+58 to +1034) and *slam*-RC (+1500 to +3427), 824.2 kb apart, *elba1*-RA (+120 to +1095) and, *slam*-RC (+1500 to +3427), 1690.7 kb apart, *CG14014*-RB (+58 to +1034) and *CG15382-RA* (+113 to +979), 3393.1 kb apart, *hb*-RA (+164to +2440) and *slam*-RC (+1500~+3427) on different chromosomes; *hb* probe was a gift from Steve Small. Gene foci pairs in a nucleus were assigned in the following manner: a focus of ‘gene A’ was assigned to the closest ‘gene B’ focus. Up to two pairs of foci can be assigned per nucleus (representing the two genes on the two homologous chromosomes). Using the Mann-Whitney Rank Sum Test for any two of the groups, pairs with distances between 1700 kb ~ 15 kb have a significant positive correlation between 1D distance and 3D distance (Figure 3H). No significant differences were noted between 3.4 Mb and 1.7 Mb pairs, or 15 kb and 4 kb pairs. A control pair of *hb* and *slam* (Figure 3G), two genes on different chromosomes, had the farthest 3D distance (2186±909 nm) and no close proximity was ever observed, as expected. Two probes against different regions of the *slam* gene (*slam* and *slam#2*, 87 bp apart) had an average distance of 185±33 nm, thus we established our resolution limit as 218 nm (mean plus one standard deviation covers 84% of data point).

### Single molecular fluorescent *in situ* hybridization (smFISH)

Antisense probes against *Bsg25D* and *CG14014* were generated following the Gasper et al. protocol [43] with minor modifications. Probes comprising 33 (Bsg25D) and 40 bp (CG14014) oligonucleotides (20 bases each; Integrated DNA Technologies) with gaps of at least two bases were NH2-dd-UTP conjugated using terminal deoxy-nucleotidyl transferase (NEB) and labeled with succinimidyl (NHS)-ester conjugated Alexa fluor 647 or Alexa flour 594 dyes (Invitrogen). smFISH was done following the Gasper et al. protocol [43]. Two different fluorophores were used for experiments comparing two different genes. For experiments combining IF and smFISH, IF was done before smFISH.

### smFISH 3D data analysis

Spots of smFISH were called the same way as Pol II spots to detect up to eight foci per nucleus, and their position and intensity were recorded. Duplicated sister chromatids were excluded from further analysis. Distances between foci of different genes were measured with the Imaris “measure” function by assigning start and end points from the center of defined foci. To compare smFISH signal of close foci (transcribed in proximity) versus distant foci (transcribed in separable distance), we chose nuclei containing one pair of close foci (<400 nm) and another pair of distant foci (>500nm) and generated their signal intensity ratio (close/distant). Comparing within each nucleus eliminates the transcriptional variance between nuclei which may enter interphase at slightly different times. We analyzed 555 nuclei in 14 embryos with this stringent criteria. We divided the close foci into three groups (<200 nm, 200-300 nm, 300-400 nm) to compare them with foci of >500 nm pairs, and their ratios are shown in Figure 4F.

