## Supplemental figures for "An “individualist” model of an active genome in a developing embryo"

### Supplemental Information

#### Figure S1

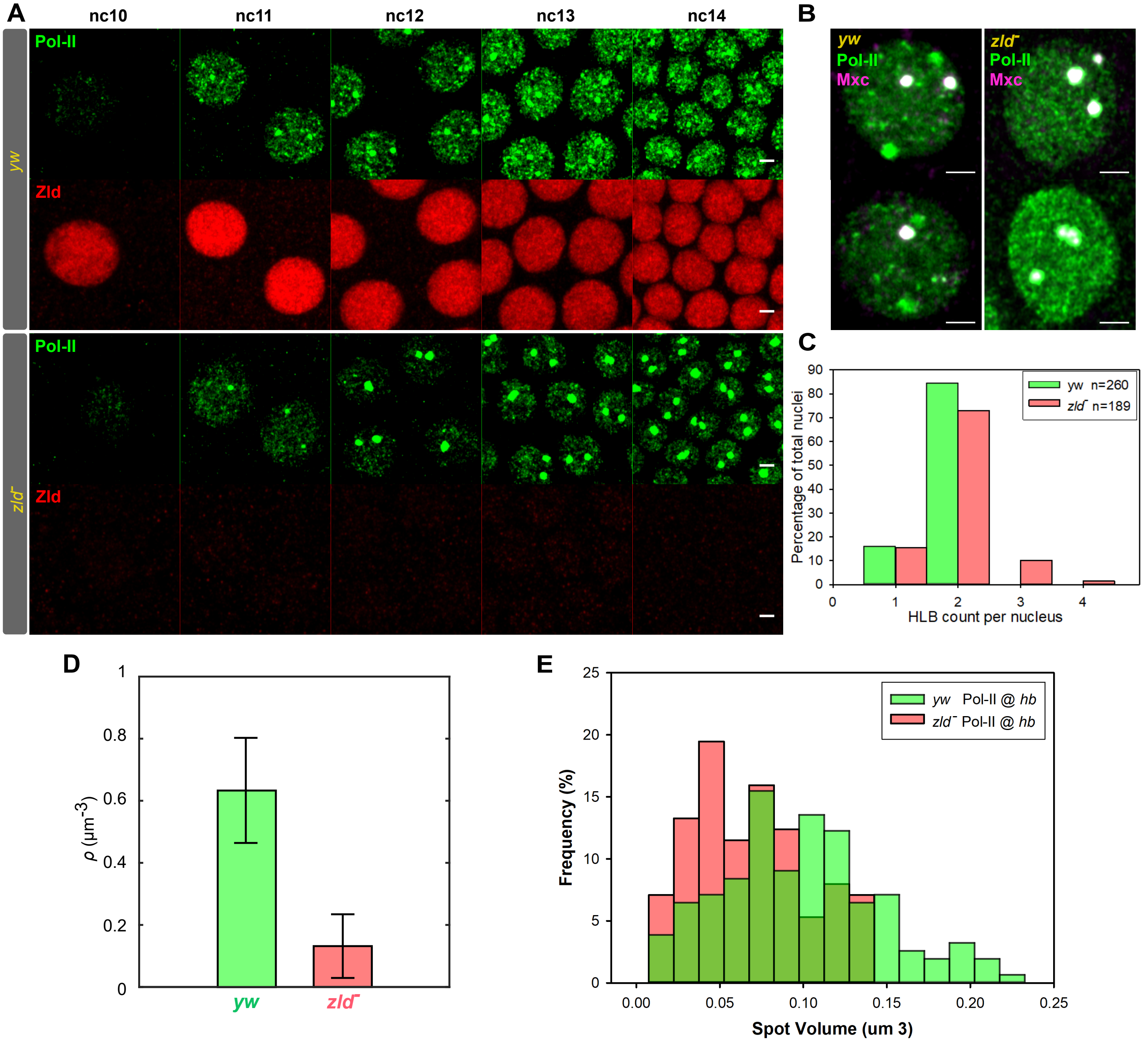


**Figure S1. Pol II form speckles at active genes and histone locus bodies (HLB).**
(A) Immunofluorescence (IF) of Pol II and Zld in *yw* and *zld*⁻ embryos which were fixed and stained together using anti-Zelda antibodies to distinguish. Scale bar = 2 μm. Note that Zelda protein is abundant and not visibly in speckles like Pol II. (B) IF of Pol II and Mxc, a HLB marker, in *yw* and *zld*⁻ nuclei. Usually one (left bottom) or two HLB were observed (left top) in *yw*. More than two HLB can only be observed in *zld*⁻ nuclei (right). Scale bar = 2 μm. (C) Statics of HLB per nucleus. 15.8% of *yw* and 15.3% *zld*⁻ nuclei have one HLB. 84.2% of *yw* and 73.0% *zld*⁻ nuclei have two HLBs.10.1% *zld* ⁻ nuclei have three HLBs. 1.6% *zld*⁻ nuclei have four or more HLBs. (D) Pol II speckle density (ρ) in *yw* (0.64±0.17 μm3) and *zld*⁻ (0.13±0.10 μm3) nuclei. (E) Distribution of volume sizes of Pol II speckles at hb loci in *yw* (n=155) and *zld*⁻ (n=113) nuclei at 75% embryonic egg length (mid anterior region of hb domain). Two embryos of each genotype were analyzed.

### Figure S2

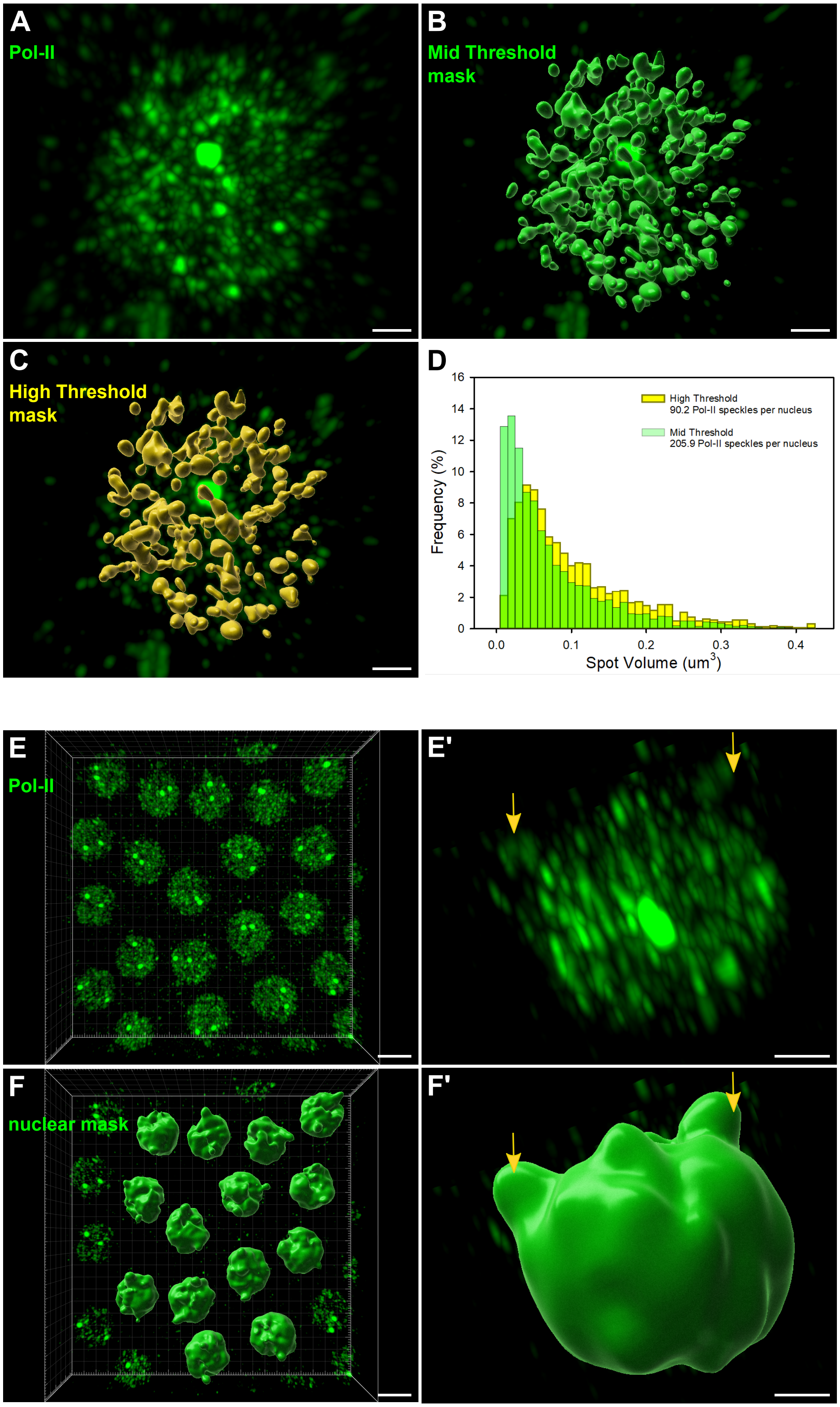


**Figure S2. Pol II speckle calling threshold setting and nuclear volume detection.** (A) Masks of 3D Pol II antibody stained images were generated by the Imaris "surface" function with either a mid-threshold of intensity (B), which included most of the spots, or a high threshold of intensity (C), which included strong spots only. Scale bar = 1 μm. (D) The distribution of volume of speckles. Note that the high threshold mask captured less tiny spots which are more likely to be noises. To determine the centers of nuclei by Imaris "surface" function, 3D Pol II antibody staining images (E) were used to generate a nuclei mask (F), then output the calculated "center of homogenous mass". Pol II antibody also detects Pol II CTD T4p in the centrosome [2] (arrow in single nucleus view of E' and F'). Scale bars are 5 μm in E and F and 2 μm in E' and F'.

### Figure S3

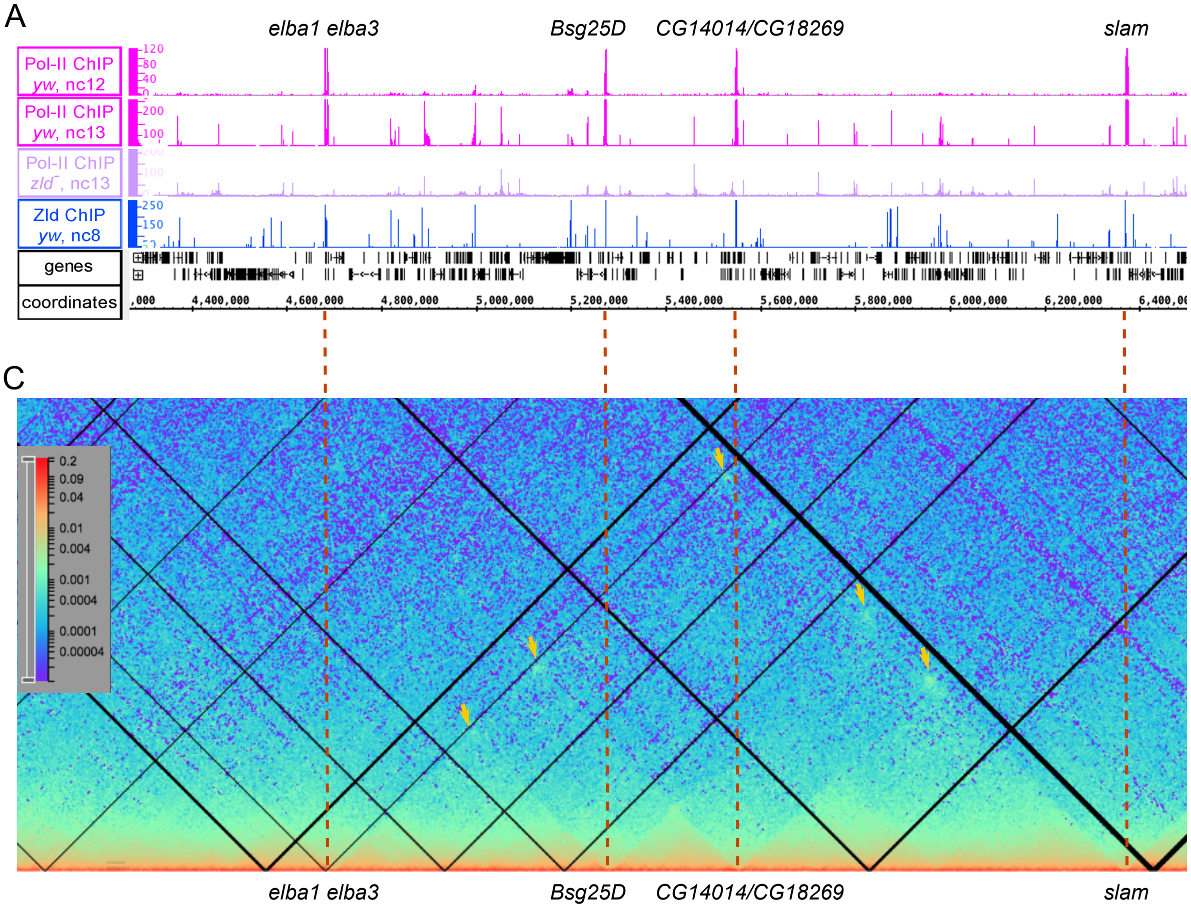


B

| | | |
|---|---|---|
| *CG7271* | 164 | 3L |
| *CR44504* | 163 | 2R |
| *bnk* | 161 | 3R |
| *slam* | 160 | 2L |
| *CG14014* | 152 | 2L |
| *elba1* | 146 | 2L |
| *CG34137* | 146 | 2L |
| *hrg* | 143 | 2R |
| *CG14317* | 142 | 3R |
| *bro* | 137 | 3L |
| *scw* | 136 | 2L |
| *kuk* | 131 | 3R |
| *elba3* | 130 | 2L |
| *CG15876* | 129 | 3L |
| *CG15382* | 128 | 2L |
| *Bsg25D* | 122 | 2L |
| *halo* | 121 | 2L |
| *sry-alpha* | 120 | 3R |
| *SNCF* | 120 | 3L |
| *Brd* | 120 | 3L |
| *tsg* | 119 | X |
| *CG34214* | 118 | 2R |
| *CG18269* | 118 | 2L |
| *nullo* | 116 | X |
| *CG8960* | 116 | 3L |
| *amos* | 114 | 2L |
| *CG43184* | 112 | 3L |
| *CG13427* | 109 | 2R |
| *elba2* | 108 | 2L |
| *CG34266* | 101 | 3L |
| *CG4570* | 100 | 3R |
| *neu2* | 100 | 3L |
| *CG13711* | 99 | 3L |
| *Taf4* | 98 | 3L |
| *CG13000* | 97 | X |
| *CG13713* | 97 | 3L |
| *CG14427* | 96 | X |
| *wech* | 96 | 2R |
| *bnb* | 95 | X |
| *CG13465* | 95 | 3L |
| *E(spl)m4-BFM* | 94 | 3R |
| *CG18132* | 94 | 2L |
| *snRNA U1:95Cb* | 93 | 3R |
| *runt* | 91 | X |
| *CG13712* | 87 | 3L |
| *CG14915* | 86 | 2L |
| *Nrt* | 85 | 3L |
| *l(1)sc* | 84 | X |

**Figure S3. TAD boundaries are near active Zelda-bound target genes.**
(A) Zelda [3] and Pol II [1] ChIP-seq tracks of a 2 Mb region on chromosome 2L. Tracks from top to bottom: Zelda in *yw* at nc8, Pol II in *yw* at nc12, *yw* at nc13, and *zld*⁻ at nc13. Note the low (nine), but highly expressed, number of genes at nc 12, the first wave of genome activation (elba1 and elba 3 are very close together and therefore appear as one peak, and the same for CG14014 and CG18269). Also note that Zelda bound peaks lie directly upstream of the transcription unit, a common feature of these genes. (B) List of the top 50 most highly Pol II bound genes, ranked by peak height at nc12. The genes shown in the ChIP-seq tracks are highlighted in purple, and other genes examined in this study are highlighted in green. (C) Hi-C data [4] of the corresponding region at nc 12 reveal boundary interactions between these genes (arrows).
